# Breast cancer risk SNPs converge on estrogen receptor binding sites commonly shared between breast tumors to locally alter estrogen signalling output

**DOI:** 10.1101/2023.10.30.564691

**Authors:** Stacey EP. Joosten, Sebastian Gregoricchio, Suzan Stelloo, Elif Yapıcı, Chia-Chi Flora Huang, Maria Donaldson Collier, Tunc Morova, Berkay Altintas, Yongsoo Kim, Sander Canisius, Gozde Korkmaz, Nathan Lack, Michiel Vermeulen, Sabine C. Linn, Wilbert Zwart

## Abstract

Estrogen Receptor alpha (ERα) is the main driver and prime drug target in luminal breast. ERα chromatin binding is extensively studied in cell lines and a limited number of human tumors, using consensi of peaks shared among samples. However, little is known about inter-tumor heterogeneity of ERα chromatin action, along with its biological implications.

Here, we use a large set of ERα ChIP-seq data from 70 ERα+ breast cancers to explore inter-patient heterogeneity in ERα DNA binding, to reveal a striking inter-tumor heterogeneity of ERα action. Interestingly, commonly-shared ERα sites showed the highest estrogen-driven enhancer activity and were most-engaged in long-range chromatin interactions. In addition, the most-commonly shared ERα-occupied enhancers were enriched for breast cancer risk SNP loci. We experimentally confirm SNVs to impact chromatin binding potential for ERα and its pioneer factor FOXA1. Finally, in the TCGA breast cancer cohort, we could confirm these variations to associate with differences in expression for the target gene. Cumulatively, we reveal a natural hierarchy of ERα-chromatin interactions in breast cancers within a highly heterogeneous inter-tumor ERα landscape, with the most-common shared regions being most active and affected by germline functional risk SNPs for breast cancer development.

## INTRODUCTION

The Estrogen Receptor alpha (ERα) is the driving force in most breast cancers diagnosed in men and women worldwide (1). As such, ERα is considered the critical drug target in both the adjuvant and metastatic phase of the disease, but resistance to hormonal treatment is common (2). ERα serves as a hormone-dependent transcription factor, associating to DNA regulatory elements upon ligand-mediated activation to drive activity of responsive genes, ultimately giving rise to tumor growth. DNA binding sites for ERα are enriched for the palindromic DNA sequence AGGTCAnnnTGACCT, termed Estrogen Response Elements (EREs), through which the ERα homodimer directly interacts with the DNA (3). To study ERα DNA action in a genome-wide and comprehensive fashion, Chromatin immunoprecipitation – followed by sequencing (ChIP-seq) was used to profile the receptor’s DNA binding pattern in breast cancer cell lines as well as in tumors. From these studies, we now know the vast majority of ERα binding sites is found further away from the genes these control (4), and around 95% of all ERα chromatin binding is found at distal intergenic regions or introns (5,6), positive for the classical enhancer marks H3K27ac and P300 (4). Interestingly, the total number of ERα sites found at putative enhancers greatly outnumbers the number of genes these controls (7), implying level of functional redundancy or cooperative action between ERα sites that is still poorly understood.

Previous studies have identified ERα binding patterns that characterize response to hormonal treatment or prognostication by sex. In cell lines, numerous studies report on plasticity in ERα DNA binding in endocrine therapy sensitive MCF7 cells, as well as their treatment resistant derivatives (6,8,9). In patients, Ross-Innes *et al.* (6) first reported on distinct ERα binding profiles and associated gene expression between patients with good (ERα+/PR+) or poor (ERα+/PR-) clinical outcome, resulting from FOXA1-mediated reprogramming. Later, our team reported an ERα ChIP-seq based classifier using primary tumor specimens, capable to identify breast cancer patient on response to aromatase inhibitor treatment in the metastatic setting (10). Inter-tumor variability in ERα DNA binding has also been described to decrease dramatically upon neoadjuvant tamoxifen treatment and associated binding sites again able to differentiate patient outcome (11). ERα DNA binding profiles were not found to differ greatly between male or female breast cancer patients, though sites associated with patient outcome were sex-specific (12). These studies highlight the value of characterizing ERα DNA binding between clinically distinguishable groups of patients. However, even within the studied groups, large variation in ERα DNA binding was observed, of which the biological and clinical implications remain unknown.

Due to the observed inter-sample heterogeneity of ERα peaks, downstream analyses typically rely on a consensus of peaks. Whether a peak is included in a consensus relies on an (often arbitrarily chosen) threshold for the minimum number of tumors in which the peak is detected. This leaves large amounts of potentially interesting data unused. To the contrary, here we set out to explore inter-patient heterogeneity in ERα binding in more detail. We evaluate the biology underlying inter-tumor cistromic heterogeneity of ERα, in relation to genomic locations and germline variations between breast cancer tumors, to better understand the possible biological implications thereof.

## MATERIAL AND METHODS

### Patient cohorts

ERα ChIP-seq on 30 male breast cancer samples was previously published by our group (12). The female cohort was compiled of previously published ERα ChIPs performed in our lab (12), 5 newly generated samples and, ERα ChIPs published by others (6,10). Only primary tumors were included. Sample details are described in Supplementary Table 1.

### Cell culture and chemicals

MCF7 human breast cancer cell lines have been cultured in Dulbecco’s Modified Eagle Medium (DMEM, Gibco) supplemented with 10% Fetal Bovine Serum (FBS-12A, Capricorn Scientific) and penicillin/streptomycin (100μg/mL, Gibco). Cell lines were subjected to regular *Mycoplasma* testing, and underwent authentication by short tandem repeat profiling (Eurofins Genomics).

For hormone stimulation, cells were pre-cultured for 3 days in phenol red-free DMEM (Gibco) supplied with 5% dextran-coated charcoal (DCC) stripped FBS, 2mM L-glutamine (Gibco), and penicillin/streptomycin (100μg/mL, Gibco), then stimulated with 10nM DMSO-solubilized 17β-estradiol (MedChemExpress, #HY-B0141) for 6 hours.

### Publicly available cell line data analysis

Called peaks of ERα ChIP-seq for MCF7 (GSM798423, GSM631484, GSM1967545), T-47D (GSE68359, GSE32222) (6,13) and ZR-75-1 (GSE25710) under full medium conditions were downloaded from Cistrome Data Browser (14,15) and subsequently lifted over from hg38 to hg19 by UCSC *LiftOver* (16). Sample details are described in Supplementary Table 1. To assess how well these cell lines represented the heterogeneous ERα cistrome found in patients, a union of cell line peaks was generated in *Diffbind* v.2.9.0. (17) and subsequently intersected with the list of patient peaks by *Bedtools* (18). The intersected peakset was subsequently used to create heatmaps shown in Supplementary Figure 1h, using *Easeq* (19).

### ChIP-seq library preparation

Five newly generated ERα samples were generated per previously published protocol (20,21). Sample details are described in Supplementary Table 1. Fresh frozen tumor material was cryosectioned, collected in Eppendorf tubes and stored at −80°C until processing. An H&E slide was assessed by pathologist to confirm tumor cell content. For ChIP, tissue was defrosted on ice and cross-linked in solution A (50 mM Hepes, 100 mM NaCl, 1 mM EDTA, 0.5 mM EGTA, pH = 7.4) containing 2 mM DSG (Sigma) and incubated at room temperature for 25 min while rotating. Next, formaldehyde was added to 1% final concentration and rotation was continued for 20 min. Reaction was quenched by 0.2 M glycine. Samples were pelleted, washed with cold PBS and tissue architecture was disrupted by a pellet pestle (Sigma) and subsequently sonicated (Diagenode PicoBioruptor). IP was performed by overnight incubation with 5 ug ERα antibody (SC-543, Santa Cruz) to 50ul dynabeads (Invitrogen) in blocking buffer (5% BSA in PBS), then washed ten times with RIPA (50 mM HEPES, 500 mM LiCl, 1mM EDTA, 1% NP-40, 0.7% Na-DOC, pH = 7.6), washed in TBS, reverse cross-linked at 65°C in Elution Buffer (50mM Tris, 10mM EDTA, 1% SDS) and DNA eluted. ChIP DNA was then prepared for Illumina multiplex-sequencing with 10 samples per lane at 65 bp single end and sequenced on Illumina HiSeq 2500 (Illumina).

### ChIP-seq data analysis

Data of all patient samples was aligned to Hg19/GRCh37 using Burrows-Wheeler Aligner (*BWA*, v0.7.5a) (22) with mapping quality above 20 and peak called with *MACS2* (v1.4) (23) using the pipeline available at https://github.com/csijcs/snakepipes. Heatmaps and peak snapshots were generated with *Easeq* (19) and all other plots in using *ggplot2* (24) under R v4.0.1. The percentage of peaks included or left out at varying thresholds (Figure 1b, d) was analyzed in *DiffBind* (17). Genomic distribution (Figure 1c) was assessed with package *ChIPpeakAnno* (v3.15.1) (25).

**Figure 1.**
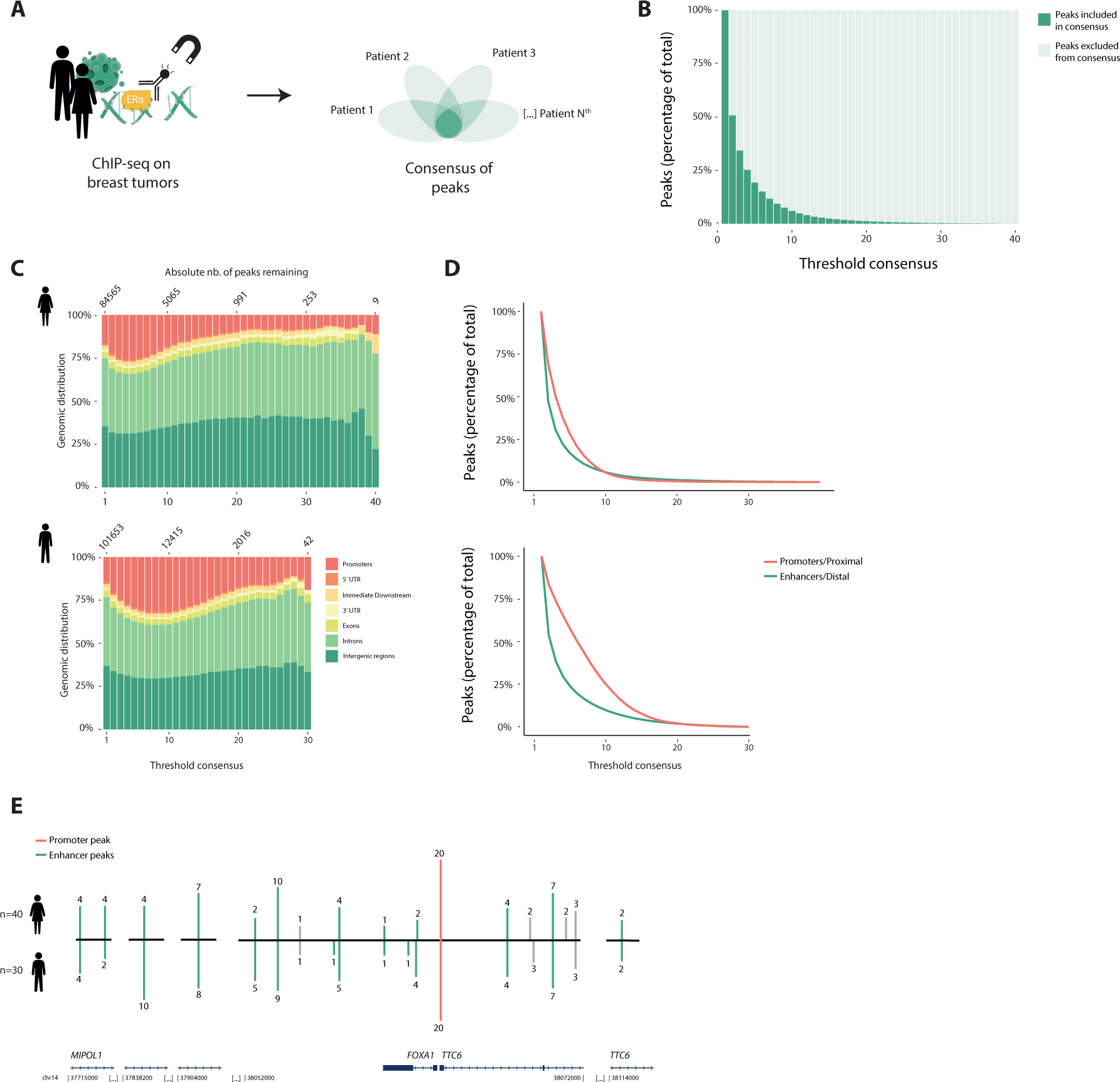
Largest inter-patient heterogeneity in ERα chromatin binding at putative enhancers. (**A**) Graphical representation of study design. ERα ChIP-seq on tumor samples from 30 male- and 40 female breast cancer patients analyzed for level of overlap and biological features. For sample details, see Supplementary Table 1. (**B**) Percentage of ERα peaks included or excluded in consensus, by varying the threshold of minimal overlap of peaks between female patients. (**C**) Genomic distribution of ERα consensus by varying threshold in females. (**D**) Percentage of distal and proximal regions retained by varying threshold for consensus in females. (**E**) ERα binding sites in the vicinity of FOXA1, showing the number of patients in which these peaks were called. Green lines represent enhancer regions, red line indicates promoter. Enhancer regions were coupled to FOXA1 on the basis of Corces *et al.*, (2018)(27). Grey lines represent peaks that were not coupled to FOXA1 on the basis of Corces et al, but these are shown for completeness as they were located in between peaks that were coupled to FOXA1.

To determine in how many patients a putative enhancer peak was called, per patient bed files were separated into promoter (defined as peak between 1-1000bp upstream of TSS, from refSeq Hg19) and putative enhancer files. A union of all patient putative enhancer peaks was generated by *Diffbind*. The union was next intersected with the per patient enhancer files again, using the option -C of *bedtools* (v2.29.2)’ *intersect* (18), producing a list discriminating between patients with and without signal for each peak in the union. Subsetting this to patients with signal and then using *bedtools intersect -C* option again produced the final list of peaks and patient counts used in this manuscript.

### Motif analysis

Presence and strength of EREs (HOMER’s MC00355) or FORKHEAD motifs (also HOMER’s) was defined by HOMER using a minimal log odds threshold of 2 (26). In case of multiple EREs or FORKHEAD motifs in a peak, the strongest was used for analyses. To assess the relationship between motif strength and heterogeneity in ERα binding, linear regression (with dummy variables) was performed in SPSS.

### Gene coupling, patterns and dependency

Breast cancer specific ATAC-seq based enhancer-promoter loops published by Corces *et al.* (2018) (27) were used to associate distal ERα binding sites to genes. Enrichment of genes patterns was assessed with GSEA 4.3.2 (28,29) For the purpose of enrichment analysis, if multiple enhancers looped to the same gene, only the binding site with the highest patient binding score was included. DepMap’s Chronos 23Q2 (Broad Institute, https://depmap.org/) dataset was used to assess relationship between commonness of ERα binding and the dependency of breast cancer cells to the associated gene.

### Hi-C library preparation and data processing

Hi-C single-index library preparation of MCF7 cells was performed as previously described using MboI (New England Biolabs) restriction enzyme (30).

Quality and quantification of the Hi-C libraries was assessed using the 2100 Bioanalyzer (Agilent, DNA 7500 kit). Four biological replicates have been pooled in an equimolar manner and subjected to sequencing using the Illumina NextSeq 550 System in a 75bp paired-end setup. De-multiplexed fastq data were analyzed at 10kb resolution using the *snHiC* pipeline (v0.2.0) (31) applying default parameters and the Hg19/GRCh37 genome assembly. Aggregate analyses at ERα binding sites have been performed using *GENOVA* (v1.0.1) (32).

### ERα-focused STARR-seq capture library design

A custom oligonucleotide probe pool (Agilent) was designed to capture ERα binding regions from clinical ChIP-seq. We selected 11,463 regions, which included all peaks that were called in at least 7 patients or more (n=7922), all regions of which coordinates intersected with rSNP coordinates (n=217), and a random sampling of less common peaks.

Pooled human genomic DNA (NA13421; Coriell Institute for Medical Research) was randomly sheared into 500-800bp fragments and ligated with Illumina compatible IDT xGen CS stubby adaptors that contain 3bp unique molecular identifiers (UMI). After the hybridizaiton of the adaptor-ligated gDNA fragments to the biotinylated probe pool, the target regions were captured with Dynabeads M-270 Streptavidin beads (Imvitrogen). The post-capture were PCR-amplified with STARR_in-fusion_Fw and STARR_in-fusion_Rv primers (5’-TAGAGCATGCACCGGACACTCTTTCCCTACACGACGCTCTTCCGATCT-3’ and 5’-GGCCGAATTCGTCGAGTGACTGGAGTTCAGACGTGTGCTCTTCCGATCT-3’), and cloned into AgeI-HF (NEB) and SalI-HF (NEB) digested hSTARR-ORI plasmid (Addgene plasmid #99296) using NEBuilder HiFi DNA Assembly Master Mix (NEB). The ERα-focused STARR-seq capture library was transformed into MegaX DH10B T1R electrocompetent cells (Thermo Fisher) and the plasmid DNA was extracted using the Qiagen Plasmid Maxi Kit.

### ERα STARR-seq library preparation analyses

MCF7 cells (>2 × 10^8^ cells/replica for 3 biological replica) were grown for 48 hrs in DMEM (low glucose, pyruvate, no glutamine, and no phenol red) (Gibco) supplemented with 5% Charcoal Striped Serum (Biowest) and 1% penicillin-streptomycin (Gibco) before being transfected with the cloned ERα-focused STARR-seq capture library using polyethylenimine (Polysciences). Following 24 hrs incubation, cells were treated with estradiol (E2) (10nM, MedChemExpress) for 6 hrs. Total RNA was extracted using TRIzol reagent (Invitrogen). The poly-A mRNA was isolated using the Oligo (dT)25 Dynabeads (Thermo Fisher), digested with Turbo DNase I (Thermo Fisher), and reverse transcribed into cDNA with the gene-specific primer (5’-CTCATCAATGTATCTTATCATGTCTG-3’). After the treatment with RNase A (Thermo Fisher), the synthesized ERα STARR-seq cDNA was amplified by a junction PCR (17 cycles) with the RNA_jPCR_Fw primer (5’-TCGTGAGGCACTGGGCAG*G*T*G*T*C-3’) and the jPCR_Rv primer (5’-CTTATCATGTCTGCTCGA*A*G*C-3’). The ERα-focused STARR-seq capture library plasmid DNA was PCR-amplified (12 cycles) with the DNA-specific junction PCR primers (DNA_jPCR_Fw, 5’-CCTTTCTCTCCACAGGT*G*T*C-3’) and jPCR_Rv primers. After purification with AmpureXP beads (Beckman Coulter), all final Illumina compatible ERα STARRseq and ERα-focused STARR-seq capture libraries were prepared by PCR amplification (7 cycles) with NEBNext universal and single indexing primers (NEB), and were sequenced on Illumina NovaSeq 6000 (150bp Paired-End).

### STARR-seq differential analyses

Raw fastq reads were mapped onto hg19/GRCh37 genome build using *BWA* (v0.7.17) aligner (22). Aligned fragments were filtered for mapping quality greater than 30 (MAPQ>30) using *samtools* (v1.14). BAM files converted into BEDPE files and fragment counts at target regions have been computed using *bedtools* (v2.30).

Differential analyses of STARR-seq data were performed using *DESeq2* (v1.30.1) (33). If fold change over DNA in both E2 and DMSO was lower than 1, the signal was considered background (*inactive*). Sites were considered *non-induced* if the linear fold change DMSO/E2 was lower than 2. If the fold change DMSO/E2 exceeded 2 and the accompanying adjusted p-value was lower than 0.00001, sites were considered as *E2-induced*.

Raw counts, *DESeq2*-normalized counts and differential analyses results are available in Supplementary Table 5.

### rSNPs analysis

Breast cancer risk SNPs accompanying Michailidou *et al.* (2017) (34) were downloaded from the Breast Cancer Association Consortium (BCAC) website (https://bcac.ccge.medschl.cam.ac.uk/). rSNP information was used from the combined Oncoarray, iCOGS GWAS meta-analysis results for ERα positive disease and only considering rSNPs with a p-value <10^−6^, excluding rSNPs that also had a significant association with ERα negative disease.

### DNA affinity purification and LC-MS analysis

MCF7 cells were harvested and washed twice with ice-cold PBS and nuclear extracts were prepared as described previously (35). ∼50bp oligonucleotide probes, encompassing the ERE roughly at the center with either WT or SNP sequence, were ordered with the forward strand containing a 5’-biotin moiety (Integrated DNA Technologies) (Supplementary Table 4). DNA affinity purifications, on-bead trypsin digestion and dimethyl labeling was performed as described (36). Matching light and medium labelled samples were then combined and analyzed using a gradient from 7 to 30% buffer B in Buffer A over 44min, followed by a further increase to 95 percent in the next 16 minutes at flow rate of 250nL/min using an Easy-nLC 1000 (Thermo Fisher Scientific) coupled online to an Orbitrap Exploris 480 (Thermo Fisher Scientific). MS1 spectra were acquired at 120,000 resolution with a scan range from 350 to 1300 m/z, normalized AGC target of 300% and maximum injection time of 20 ms. The top 20 most intense ions with a charge state 2-6 from each MS1 scan were selected for fragmentation by HCD. MS2 resolution was set at 15,000 with a normalized AGC target of 75%. Raw MS spectra were analyzed using MaxQuant software (version 1.6.0.1) with standard settings, with multiplicity set to 2 with dimethyl Lys 0 and N-term 0 as light labels and dimethyl Lys 4 and N as medium labels and requantify enabled (36,37). Data was searched against the human UniProt database (fasta file downloaded 201706) using the integrated search engine.

## RESULTS

### Putative enhancers represent the largest source of inter-patient heterogeneity in ERα chromatin interactions

To identify the level of ERα binding heterogeneity in human breast cancer specimens, we used ERα ChIP-seq data from 40 female and 30 male breast cancer patients (Figure 1a, Supplementary Table 1). Five newly generated ERα ChIPs on female tumors were added to this study. The remaining 35 female and 30 male samples have been described and analyzed in previous publications, to identify genomic regions that could stratify patients on outcome or sex (6,10,12) (Supplementary Table 1). All samples were processed with the same bioinformatics pipeline (see methods for details).

Analyzing these samples, we observed a high level of inter-tumor heterogeneity of ERα binding, with the vast majority of ERα sites being poorly conserved between tumors from female patients (Figure 1b). For instance, if we were to re-evaluate the aforementioned data by consensus with a lenient threshold of peaks present in at least 2 patients, 50% of data would be ignored. Previous analyses have been performed with cut-offs as stringent as peaks found in 75% of patients (6). Applying this cut-off to this cohort, merely 0.3% (for female tumors) and 1.1% (for male tumors) of sites would remain. Both in female and male breast cancer patients, the level of conservation for ERα sites depends on the genomic distribution of the peakset (Figure 1c), as promoter binding events are more conserved between individual tumors than distal ERα binding (Figure 1d).

For both sexes, we mapped all ERα binding events to genes using promotor-enhancers loops defined by Corces *et al.* (27). Interestingly, the gene with the most distal ERα binding events identified in our patient cohorts – thus the most heterogeneous – was associated with FOXA1. FOXA1 is the classical pioneer factor essential for ERα to facilitate its chromatin binding (8). In vicinity of a homogenously-bound promoter, we found the adjoining region to be frequented by ERα at numerous putative regulatory elements and their occupancy varying strongly between patients (Figure 1e).

### Common peaks represent 30% of ERα binding in a patient ChIP sample

To better appreciate the functional implications of enhancer heterogeneity among patients, we removed all ERα binding events at promoters and ranked all distal ERα binding events (74,438 in females, 91,712 in males) from common among patients to patient unique events (Figure 2a, Supplementary Figure 1a, Supplementary Table 2, −3). Strikingly, 53% of all ERα-bound putative enhancers found in female tumors were patient unique, 46% of sites were found in 2 to 19 female patients and merely 1.1% were bound in more than half of the female cohort (Figure 2a). Similarly, among males, inter-patient heterogeneity was large with 46.3% of distal ERα binding events being patient unique, 49.5% found in 2 to 14 patients and merely 4.3% found in more than half of all male tumors analyzed (Supplementary Figure 1a).

**Figure 2.**
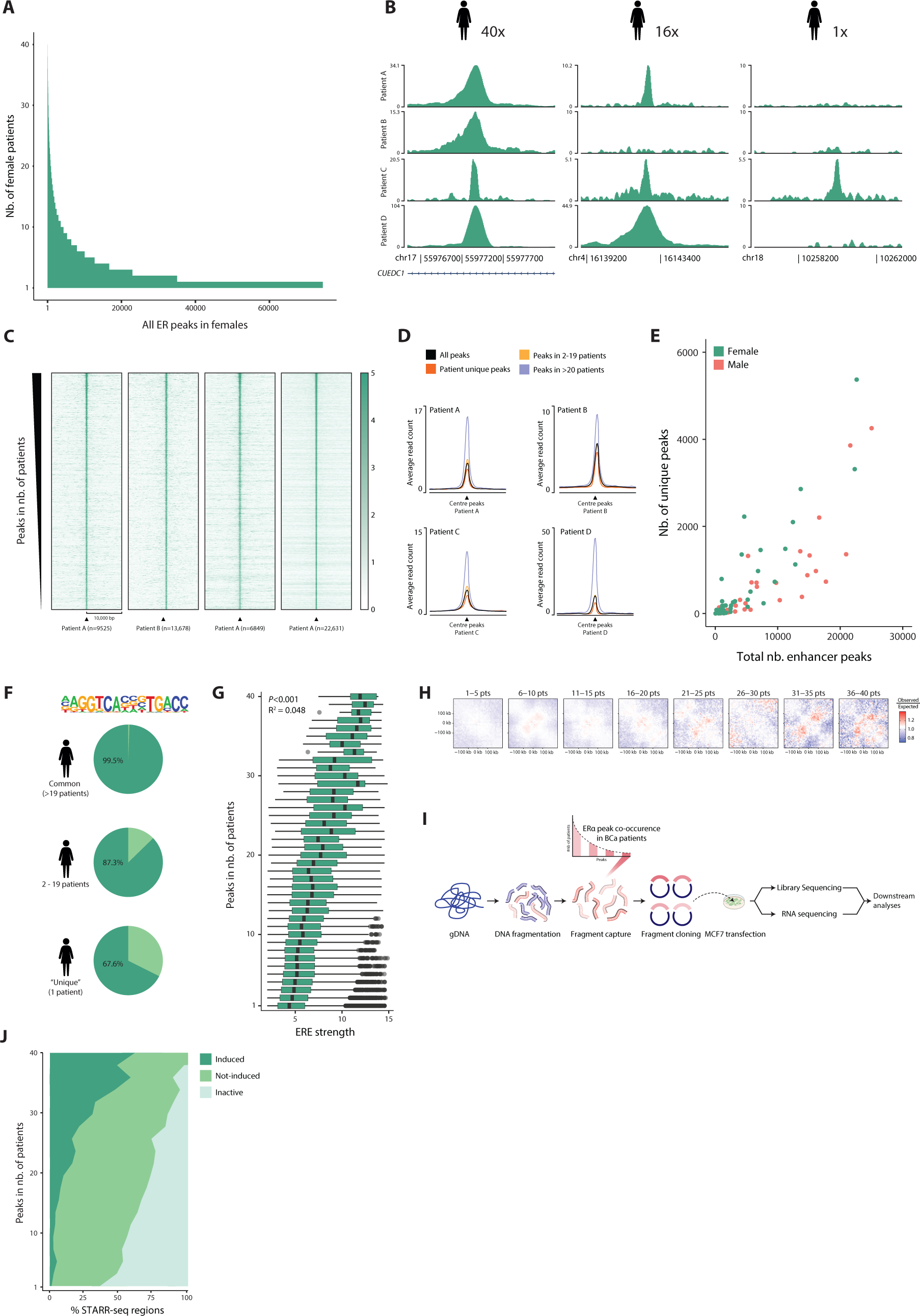
Characterization of enhancers ranked from commonly to less frequently bound by ERα shows distinct biological features. (**A**) A ranked overview of 74,438 distal ERα peaks showing in how many tumor samples each peak was found, in a cohort of 40 female patients. (**B**) Examples of ERα peaks that were peak called in tumor samples in all 40 females (left), in 16 females (middle), as well as in only one female patient (right). (**C**) Examples of per patient heatmaps of ERα signal, of peaks called in that female patient sample, ranked as in a. (**D**) For more commonly occurring and unique peaks, examples of the average intensity of ERα ChIP-seq signal in 4 female patients are shown. (**E**) Correlation plot of the total number of distal peaks in a patient sample (x-axis), versus the percentage of patient unique peaks in that sample (y-axis). (**F**) The percentage of common, less common and patient-unique ERα peaks in females that contain an estrogen response element (ERE). (**G**) The strength of those EREs as determined by HOMER, ranked from those in common to those in more patient unique ERα peaks. Black dots represent outliers. (**H**) Aggregate Region Analyses (ARA) showing the average Hi-C contacts (observed over expected scores) at ERα binding sites shared by an increasing number of patients from left to right. The matrices include a window of ±250 kb from the ERα peak centers. (**I**) Schematic overview of STARR-seq methodology. (**J**) Distribution of enhancer activity as determined by STARR-seq upon 6h of 10nM estradiol stimulation, from common to more patient unique peaks. Details on cut-offs for categories induced, not-induced and in-active are described in the Methods section.

Differences in peak conservation between patients was not due to subtle differences in peak calling performance, as partially shared ERα sites were genuinely differential enriched between tumors (Figure 2b). On patient level, signal intensity at ERα sites is higher at common as opposed to patient-unique peaks (Figure 2c, -d), with highly common peaks (>19 patients) showing strongest signal. Only 8 peaks were found conserved between all female patients and 34 in all male patients. A few well known-estrogen-related genes are proximal to top common peaks, including PGR, RARA, IGFBP4, CUEDC1 and GREB1 (Supplementary Table 2). Common peaks relate to canonical early and late estrogen response genes more frequently (Supplementary Figure 1b), although less common and unique peaks were also associated to classical estrogen-related signalling genes such as CCND1 and TFF1 (Supplementary Table 2).

For each patient in our cohort, we looked into the amount of ERα ChIP-seq peaks picked up in their respective tissue sample (Supplementary Figure 1c). When viewing the list of all distal ERα ChIP-seq peaks found in a single patient, the minority of that list is made up of peaks that we consider common among individuals in the cohort: ±29.3% (in females) and ±42% in males (Supplementary Figure 1d). This implies that the body of ERα DNA binding events in a single patient sample happen at regions considered less common among individuals. Thus, while stringent consensus-based analyses focus on peaks common among individuals, they do not take into account most ERα binding in a single individual. Or in other words, a consensus analyzes ERα on group level, but may inform less on ERα behavior in an individual patient.

In an individual patient, peaks that are unique to that patient make up 13.3% (in females) and 8.9% (in males) of all distal ERα peaks found in that tissue sample (Supplementary Figure 1d). While generally weaker, signal at patient unique sites is only slightly lower than average signal for that particular tumor (Figure 2d). The number of patient-unique peaks in a patient sample increases with the total number of distal peaks picked up in that patient sample (Figure 2e, Supplementary Figure 1c, -d), implying unique peaks are not a proxy of relatively poor ChIP-seq quality.

Cumulatively, these data confirm a remarkable inter-patient heterogeneity of ERα enhancer action, that is typically overlooked in consensus-based analyses.

### ERE strength weakly associates with the level of inter-tumor conservation of ERα sites

From cell lines, ERα DNA binding is known to be enriched at EREs, though indirect chromatin associations by tethering though other transcription factors may also occur (3). In our ranked list of peaks in female tumors, an ERE could be found in 99.5% of common peaks (>19 patients), in 87.3% of peaks found in 2-19 patients and in 67.6% of patient unique peaks (Figure 2f). We used HOMER (26) to determine the strength the EREs – that is, sequences most similar to the consensus ERE receive a higher score versus those deviating from the consensus ERE. When present, the average strength of the ERE was found to correlate with how often a binding site was bound by ERα in our cohort of female patients (*p*<0.001, Figure 2g). Nonetheless, variance in ERE strength per patient number is large and strong EREs were still observed at less common sites and vice versa. In a regression model where strength of the ERE in a peak was evaluated as predictor for the number of patients in which a ERα□peak was found, *R^2^* = 0.048 (Goodness of fit) was only 0.048, suggesting that while ERE strength is statistically significantly associated with ERα site conservation, it is not a powerful predictor of heterogeneity in ERα binding.

ERα does not act independantly, but requires activity of other proteins for its function. One essential ERα interactor is the forkhead protein FOXA1, which serves as a pioneer factor rendering the chromatin accessible for ERα to bind (8). The consensus FOXA1 motif TGTTTAC is generally found close to ERE sequences, yet does not overlap (38). In our female cohort, 87.1% of common ERα peaks, 79.6% of peaks in 2-19 patients and 62.1% of patient unique peaks carried a forkhead motif (*p*<0.001) (Supplementary Figure 1e), but no relationship between strength of the forkhead motif and heterogeneity in ERα DNA binding was observed (Supplementary Figure 1f).

### Estrogen-induced enhancer activity is highest for commonly shared ERα binding sites

We next questioned if the ERα binding profiles found in the most commonly used ERα+ breast cancer cell lines MCF7, T47D and ZR-75-1 are representative of the ERα DNA binding identified in primary human tumors. Comparing ERα□peaks from these cell lines in full medium conditions to peaks found in patients, we found that these cell lines capture 99.7% of peaks present in 19 patients or more, and 65.3% of peaks present in at least 2 to 19 patients (Supplementary Figure 1g, -h). 29% of the “patient unique” peaks were also called in at least one cell line, further solidifying the confidence in the ERα signal at these locations. ERα ChIP-seq from MCF7, T-47D and ZR-75-1 appears to recapitulate a significant number of ERα peaks found among 40 patients and we therefore deemed these cell lines adequate models to investigate functional consequences of ERα enhancer heterogeneity.

Using one of these cell line models, MCF7, we further investigate the biological features at commonly and less frequently bound ERα sites. For this, we first evaluated chromatin conformation behavior of common and more patient-unique sites by means of high-throughput chromatin conformation analyses (Hi-C), which illustrated a direct positive associated of long-range chromatin interaction frequency with the level of ERα site conservation among patients (Figure 2h). While classical ERα target genes were represented among the most-commonly shared regions (Supplementary Table 2), no enrichment for essentiality was found for genes proximal to ERα sites (39,40), relative to the level of inter-tumor heterogeneity (Supplementary Figure 1i).

We then surveyed enhancer activity across the ranked peaks using self-transcribing active regulatory region sequencing (STARR-seq) (41) (Figure 2i). This method allows for the massive parallel testing of intrinsic enhancer activity of DNA fragments, by cloning these sequences downstream of a core promoter and then quantifying the enhancer activity based on the self-transcription in mRNA transcripts. We generated a library of 11,147 regions, which included 7922 peaks that were called in at least 7 patients or more and a random sampling of less common peaks. The library was transfected into MCF7 cells and reporter read-out was generated under estradiol (E2) stimulation or vehicle control. Out of all 11,147 ERα sites cloned in the library, 597 (5.2%) were found E2-induced, 5777 (51.8%) were not-induced and 5053 (44.1%) were inactive (Supplementary Figure 1j) (See Methods for details on these categories). Of note, these distributions of observed activities were comparable to that of androgen receptor (AR) sites studied in LNCaP prostate cancer cells (42) or glucocorticoid receptor (GR) sites studied in lung cancer A549 cells (43). Overall this suggest that only a small fraction of nuclear receptor sites are actively engaged in transcriptional regulation. By plotting the peaks from common to less frequently bound by ERα in patients, we observed that these binding sites have distinct intrinsic properties. Interestingly, the more common ERα peaks among patients, the higher the percentage of estradiol-induced enhancer activity (Figure 2j). Enhancer activity at patient unique sites is slightly more often constitutively active, acting in an ERα-independent manner, though the receptor does bind.

Cumulatively, these results imply direct biological consequences of the observed inter-patient heterogeneity of ERα, with enhancers showing hormone-induced activity being mostly conserved among patients.

### Breast cancer risk SNP loci are enriched at ERα sites commonly shared by tumors

Though we observed a relationship between average ERE strength and commonness of ERα binding to enhancers among patients, this was not sufficient to explain the large observed variation. We therefore hypothesized genetic variation at enhancer elements contributes to this inter-patient heterogeneity of ERα enhancer action. To test this hypothesis, we turned to a known source in variation of breast cancer risk and analyzed our data for possible overlap with breast cancer risk single nucleotide polymorphisms (rSNPs) and small indels. Importantly, rSNPs for different cancer types have been found previously to be enriched in enhancer regions, with prostate cancer risk SNPs found enriched at AR sites (44) but also breast cancer rSNPs have been found enriched at ERα-bound regulatory elements (45–47). Any association of rSNPs or small indels with ERα site heterogeneity between tumors, remains unexplored.

We hypothesized that alterations in DNA sequence by rSNPs or small indels may affect ERα binding and thereby facilitate inter-patient heterogeneity. rSNPs and indels with significant (p<10^−6^) correlation with ERα+ breast cancer risk as published by Michailidou *et al.* (2017) (34) were tested for overlap with ERα binding sites in our cohort, yielding a combined list of 318 rSNPs and a small number of indels that overlap with ERα sites identified in our patient samples (Figure 3a). Surprisingly, the rSNPs and indels showed relative enrichment at the top of the ranked peaks, converging more often at ERα binding sites common between patients (Figure 3b, -c; *p*<0.0001). The relative enrichment of rSNPs and some indels at common ERα binding sites was also seen in the male breast cancer samples (Supplementary Figure 2a, -b; *p*<0.001). When normalized to number of bases in peaks (to exclude enrichment being an artefact of peak width, as genomic location of common sites may be slightly broader due to merging of more info) (Figure 3c and Supplementary Figure 2b) or when leaving out patient unique peaks (data not shown), the statistically significant association holds. Interestingly, in both sexes, the more commonly bound an ERα site with which a rSNP/indels coordinate(s) overlaps, the stronger the p-value for the association with breast cancer for that rSNP/Indel (Figure 3d and Supplementary Figure 2c), though the protective (negative beta) or risk effects (positive beta) of the rSNPs/indels at common sites are modest (Figure 3e).

**Figure 3.**
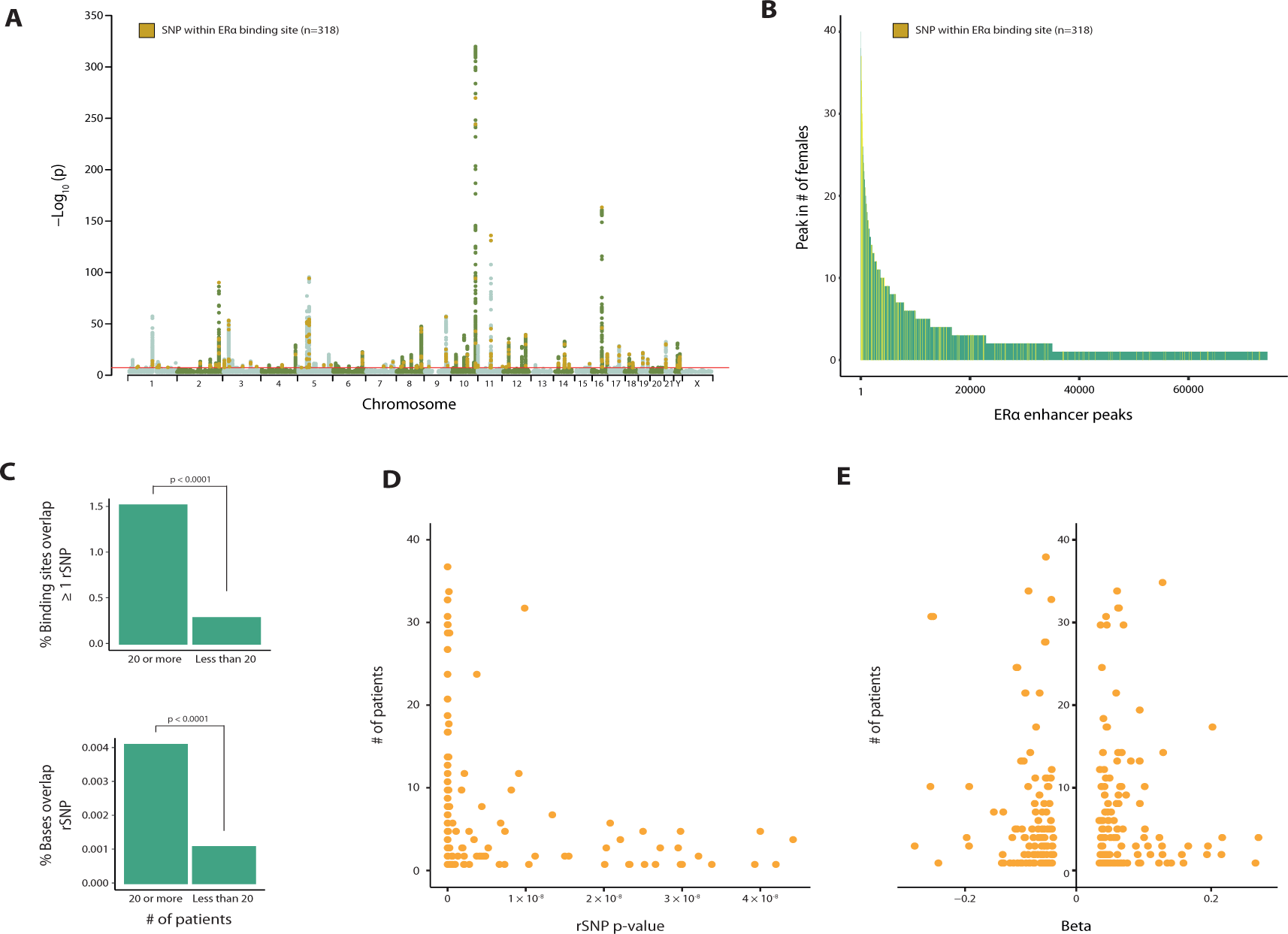
ERα+ breast cancer rSNPs enrich at regions with low inter-patient heterogeneity in Erα. (**A**) Manhattan plot of ERα+ breast cancer risk SNPs (rSNPs) with genome-wide significance originating from Michalidou *et al.* (2017) (34). Highlighted in orange are 318 rSNPs, of which the coordinates intersect with one of the 74,438 ERα peaks found among 40 female breast cancer patients. (**B**) The position of these 318 rSNPs in the ranked peaks introduced in Figure 2a. (**C**) *Top*: Comparison (Fischer exact test) of percentage of ERα *peaks* of which coordinates overlap with at least once rSNP coordinate, for common and less common ERα peaks. *Bottom*: Comparison (Fischer exact test) of the percentage of *bases* present in common or less common ERα peaks, that overlap with at least once rSNP coordinate. Even when excluding patient unique peaks, p-values remain statistically significant (data not shown). (**D**) Correlation between p-value of rSNP (x-axis) and its position in the ranking of ERα peaks introduced in Figure 2a (y-axis). If multiple rSNPs overlapped the same ERα peak, the strongest p-value was used for analysis. (**E**) Overview of beta values corresponding to rSNPs of which coordinate intersected an ERE. Negative beta values correspond with rSNPs that confer less risk to breast cancer, while positive beta values correspond to increased risk of ERα+ breast cancer.

In accordance with literature showing that rSNPs often lie in regions with transcription factor motifs (45–47), ERα peaks that overlap with rSNP/indel coordinates do hold an ERE more often than those that do not overlap with rSNP/indel coordinates (Supplementary Figure 2d). These EREs also tend to be slightly stronger than EREs without rSNPs/indels (*p*=0.06) (Supplementary Figure 2e). Though ERE strength was a statistically significant but not a powerful predictor of observed heterogeneity in ERα binding, we nonetheless checked if the enrichment of rSNPs/indels at common peaks was confounded by ERE strength, but we found no evidence to this (Supplementary Figure 2f).

### Breast cancer risk SNPs affect ERα inter-patient heterogeneity through TF motif perturbation, with biological implications on gene expression

Of the 318 rSNP/indel coordinates overlapping with ERα peaks in our cohort, 25 of those rSNP/indel coordinates actually fall within an ERE palindromic sequence (Supplementary Table 4). Of those, we focused our attention on rSNPs/indels in sites that showed enhancer activity in the STARR-seq data (E2-induced, constitutively active or not induced) (Supplementary Figure 2g). To determine which remaining rSNPs/indels had the potential to significantly and directly affect the strength of the ERE and thereby ERα binding, we used the *in silico* prediction tool *SNP2TFBS* (48), that compares the position weight matrix of a transcription factor binding motif in reference format and when including the rSNP/indel variant. This resulted in a shortlist of 3 rSNPs rs9952980, rs11695384 and rs11665924, which *SNP2TFBS* predicted to affect a hormone response element (Supplementary Table 4).

To validate the effects of these rSNPs on ERα binding experimentally, we designed two 50bp oligonucleotides containing the WT and rSNP-affected ERE, which were both biotin-labeled, and pulled through an MCF7 lysate to detect interacting proteins via mass-spectrometry in an unbiased fashion (Supplementary Table 4) (35). For rs11695384 and rs11665924, we were unable to confirm differential binding of ERα, leaving rs9952980 for further study.

Rs9952980 is a rSNP located in an intron of the gene *SLC14A2*, located on chromosome 18. This region was bound by ERα in 11 female (examples in Figure 4a) and 6 male -tumor samples. STARR-seq data confirmed the region to be active and induced upon E2 treatment (Supplementary Figure 2g). Considering the reference genome, the region holds a relatively strong ERE at a log odds motif score of 12.3. Rs9952980 affects the fifth nucleotide of the ERE (Figure 4b), which is a position of high importance for strong ERα affinity (Figure 2f), as it facilitates direct DNA-protein interaction with ERα (3). Accordingly, *in silico* analysis using *SNP2TFBS* (48) predicted Rs9952980 to significantly *decrease* ERα’s binding affinity (Figure 4c, Supplementary Table 4). Mass-spectrometry (Figure 4d) and western blot (Figure 4e) of DNA-oligo pulled-down proteins confirmed diminished ERα binding in the rSNP condition. Clinically, carriers of the alternative allele (T) have less risk (beta: −0.0549) of developing breast cancer than homozygous carriers of the reference allele (C). rs9952980 was previously predicted to regulate expression of its target gene *SLC14A2* (47) and in TCGA data, we indeed find rs9952980 significantly associates (*p*=0.0063) with reduced expression of *SLC14A2* (Figure 4f), likely mediated via the rSNPs direct impact on ERα-DNA binding.

**Figure 4.**
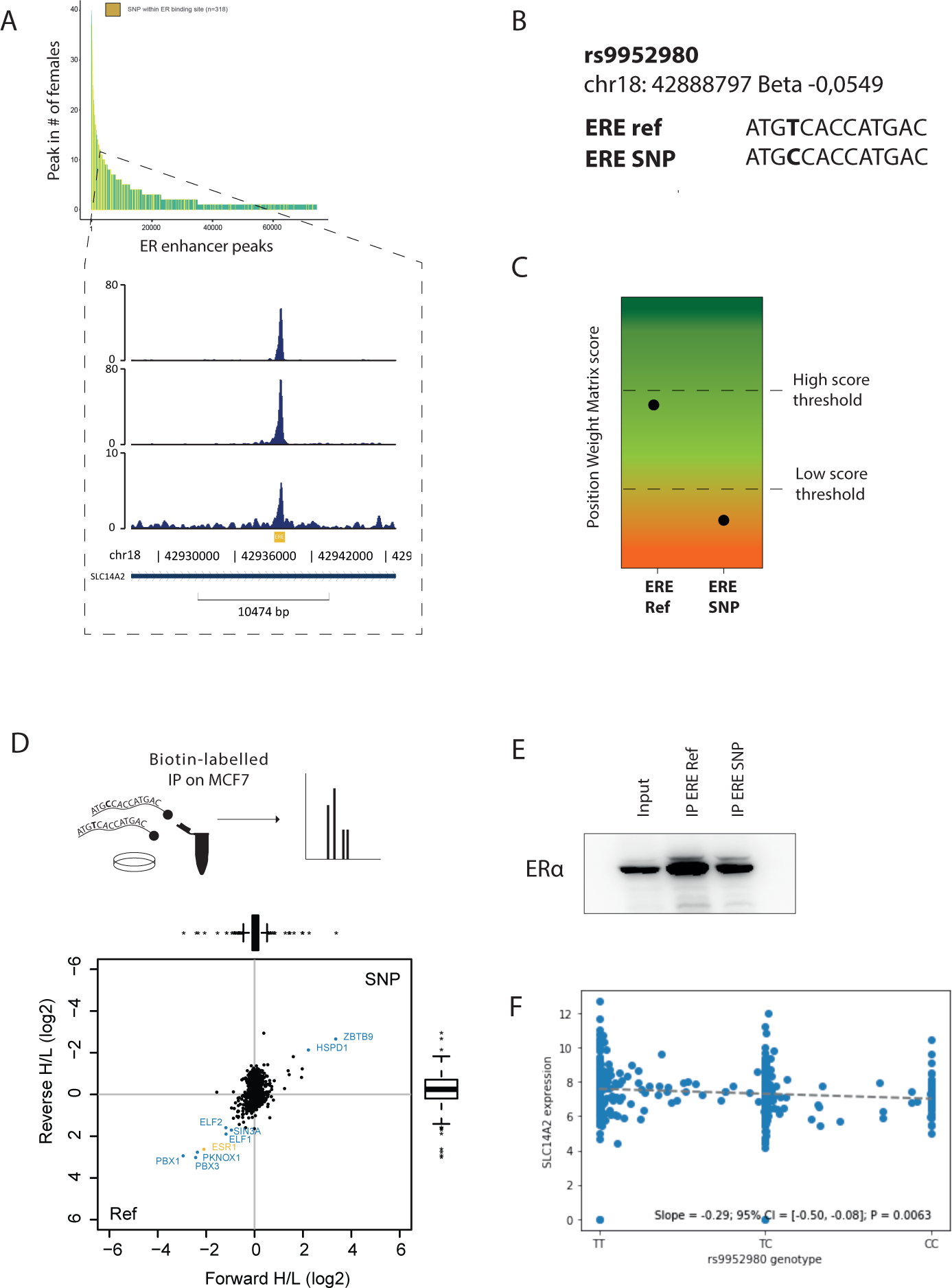
rs9952980 affects *SLC14A* expression via reduced ERα binding by impacting ERE. (**A**) Snapshots of ERα peak intersecting the coordinate of rs9952980. The peak, positioned in an intron of *SLC14A2*, was found in 11 female patients. (**B**) Estrogen response element (ERE) at this peak, in reference allele and rSNP format. (**C**) Predicted score of position weight matrix for WT and rSNP ERE, by *SNP2TFBS* (48). (**D**) Using MCF7 lysate, an immunoprecipitation (IP) was performed with 50bp biotin-labelled oligos containing the WT or the rs9952980 variant of the ERE, followed by mass-spectrometry. (**E**) ERα Western Blot (WB) of IP by 50bp biotin-labelled oligos containing the reference allele or the rs9952980 variant of the ERE. (**F**) TCGA Gene expression of *SLC14A*, which rs9952980 is predicted to affect, by homozygous or heterozygous genotype.

Few rSNPs/indels that intersected with our ranked peaks directly overlapped an ERE, though were often located in close proximity. Such rSNPs/indels may affect ERα binding indirectly, by affecting affinity of ERα’s partners, such as FOXA1. An example for this is found for rSNP rs6420415, located in an intron of *CDYL2* and of which the region was occupied by ERα in only 4 females (Figure 5a) and 4 males. rs6420415 is predicted to perturb the FORKHEAD motif such that FOXA1 binding is negatively affected (Figure 5b, -c). Indeed, western blot of oligo-mediated immunoprecipitation of the local forkhead motif in reference allele and rSNP format confirmed diminished FOXA1 binding (Figure 5d). Coinciding with these *in vitro* data, in a female tumor sample for which both ERα and FOXA1 ChIP-seq data were available, we noted ±half of the reads from the reference allele (T) and 50% from the SNP allele (G) in the ERα ChIP, while reads from the FOXA1 ChIP-seq were dominated by the reference allele T (Figure 5e). Carriers of rs6420415’s G allele are thought to have an elevated risk of developing breast cancer (beta: 0.0682) (34), possibly mediated via reduced expression of *CDYL2* (Figure 5f). *CDYL2* has been described to exert both tumor suppressing and oncogenic effects, depending on its isoform (49,50), though this distinction was not made in GWAS studies (34).

**Figure 5.**
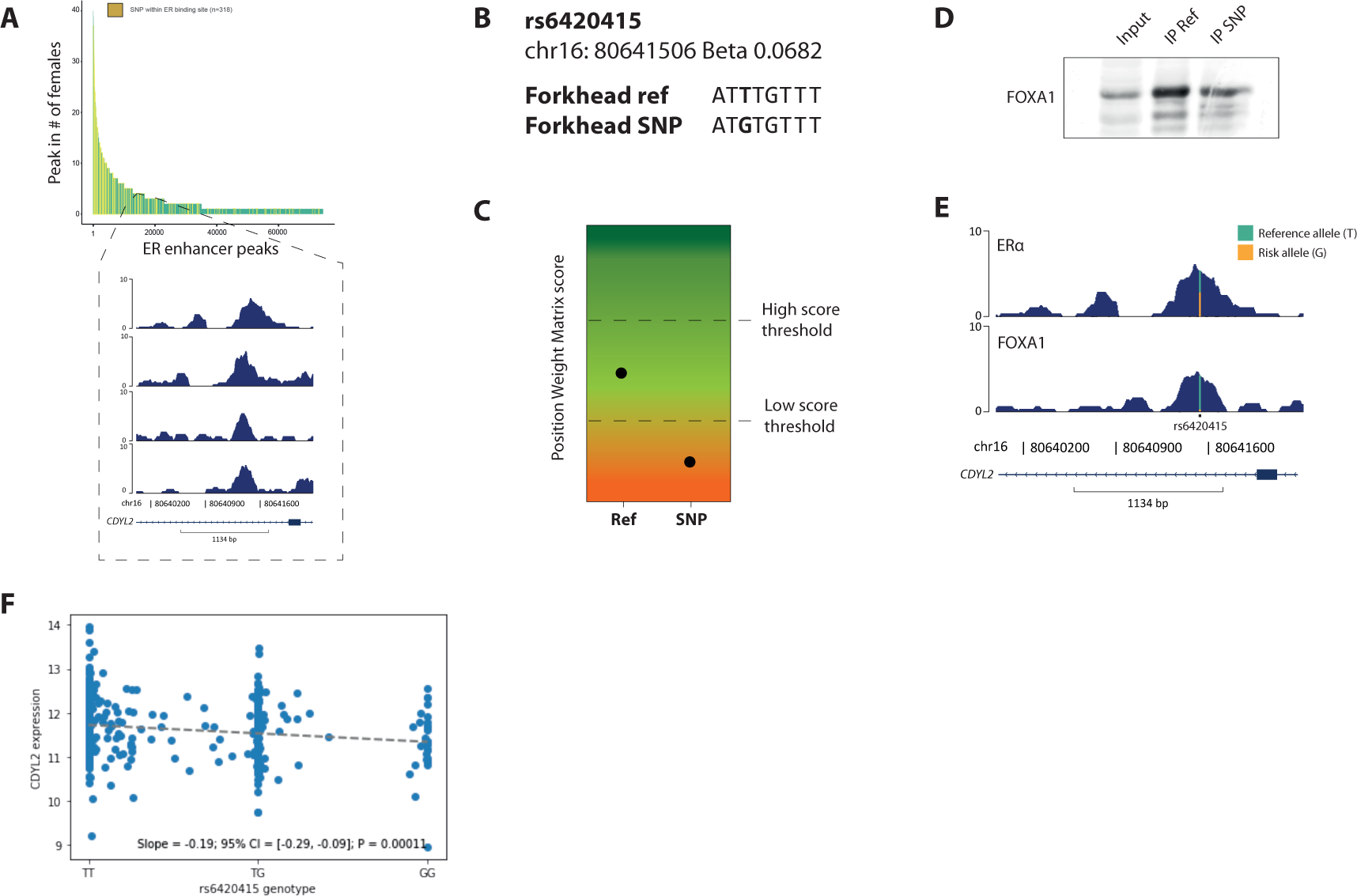
rs6420415 affects *CDYL2* expression via reduced FOXA1 binding by impacting FORKHEAD motif. (**A**) Snapshots of ERα peak intersecting the coordinate of rs6420415. The peak, positioned in an intron of *CDYL2*, was found in 4 female patients. (**B**) Forkhead motif at this peak, in reference allele and rSNP format. (**C**) Predicted score of position weight matrix for reference allele and rSNP Forkhead motif by *SNP2TFBS* (48). (**D**) FOXA1 Western blot of pulldown by 50bp biotin-labelled oligos containing the WT or rs6420415 variant of the forkhead motif. (**E**) Distribution of reads in the ERα- and FOXA1 ChIP-seq peak performed on tumor tissue from the same breast cancer patient, at the locus surrounding rs6420415. (**F**) TCGA gene expression of *CDYL2*, which rs6420415 is predicted to affect, by homozygous or heterozygous genotype for rs6420415.

Nonetheless, our data cumulatively illustrate that breast cancer risk SNPs/indels are enriched at commonly-shared active enhancer elements, can perturb binding of ERα or its pioneer factor FOXA1, for instance via allele preferential binding, which is associated with affected expression of the genes they control.

## DISCUSSION

Breast cancer is a heterogenous disease, with clearly distinct inter-tumor differences on subtype, aggressiveness and ultimately patient prognostication. Here, we show that on epigenetic scale, substantial inter-patient heterogeneity of ERα chromatin binding capacity is found. Therefore, consensus-based analyses of ERα ChIP-seq on patient samples, a strategy often applied in the field to reduce data complexity and limit noise, but at the same time eliminates potentially interesting and biologically meaningful data. Here, we queried the full spectrum of inter-patient ERα enhancer heterogeneity by ranking all peaks in patients from commonly shared to patient unique. Many peaks could only be found in a handful of patients, and around half of all peaks identified, were patient unique. As this level of heterogeneity was substantially higher for putative enhancer elements as opposed to promoters, our data suggest a level of functional redundancy between enhancers in regulating the same gene, as was recently shown in combinatorial CRISPR screening analyses (51).

While extensive QC analyses were performed on the ChIP-seq datasets, we cannot exclude that a fraction of the ERα heterogeneity between patients in our cohort may be of technical origin. The female samples in this cohort were produced in different labs, but a single antibody was used. Also, a similar degree of interpatient-heterogeneity was seen in the male cohort, of which ChIP-samples were exclusively produced in our lab, with the same (batch of) antibody. Interestingly, while inter-patient heterogeneity of ERα signal was high, our analyses on ERE strength, enhancer activity (STARR-seq data) and rSNP/indel analyses suggest that most functional activity, hormone-induced action and clinically-relevant information is found at the commonly-shared ERα sites. Thus, based on these observations, we conclude the observed enhancer heterogeneity represents a biological hierarchy of ERα action, with biological and clinical consequences.

Following this hierarchy, we found rSNP/indel coordinates to be enriched at most-commonly shared ERα peaks, in both a cohort of 40 female as well as 30 male samples. How often ERα binds to DNA at these regions in patients has never been reported before. Yet peculiarly, GWAS p-values corresponding to these rSNPs/indels are stronger at more common ERα sites, while corresponding beta values were relatively modest. One could be tempted to interpret these rSNPs/indels as statistically significant but biologically unimportant, but this does not reconcile with their enrichment at common ERα sites Hodge *et al.* (2017) (52) have hypothesized that rSNPs with strong p-values and small effects sizes may actually result from heterogeneity in the studied cohort. In this case, breast cancer risk may indeed be caused by a large number of variants and associated genes (i.e. polygenic risk model), but rather than each locus contributing a small amount to breast cancer risk, loci commonly bound by ERα contribute major risks to *distinct* breast cancers. Indeed, the patients in our cohorts have been described to have different prognoses, despite all having ERα breast cancer. When bound by ERα in almost all tumors, this also allows common sites to contribute differently to different subtypes of ERα breast cancer and thereby cause an attenuation in effect size in GWAS studies. Neither in Michalidou *et al.* (2017) (34) and nor in this work, a distinction in subtypes of ERα breast cancer was not made. To test this hypothesis, follow-up studies are required.

Cumulatively, these findings contribute to our basic understanding how sequence variants at specific regulatory elements contribute to ERα+ breast cancer development. Recently, we reported a comparable observation in prostate cancer when analyzing inter-tumor heterogeneity of Androgen Receptor action between primary tumors, revealing not only somatic mutations but also rSNPs being enriched at more-commonly shared regions, that were more active on transcriptional level (42). In that setting, we also observed that less-commonly shared regions were associated with disease progression, and may become engaged later on, at the metastatic disease stage. Future studies should address whether this phenomenon would also occur in breast cancer, and whether selectivity and plasticity of enhancer action is a more general feature in hormone-driven cancers.

In conclusion, our analyses suggest a hierarchy of ERα-chromatin interactions in breast cancers, resulting in a high degree of inter-patient heterogeneity in ERα enhancer action. We find most-commonly shared ERα regions as most-hormone inducible enhancers and serving as hotspots for germline functional risk SNPs/indels for ERα+ breast cancer development, highlighting a new perspective in better understanding the biological basis of risk variants in breast cancer.

## DATA AVAILABILITY

ERα ChIP-seq on 30 male breast cancer samples are available at Gene Expression Omnibus (GEO) (53) under accession number GSE104399 (12). The female cohort raw data can be found at GEO under accession numbers GSE104399 (12), GSE32222 (6) and GSE40867 (10). Raw data corresponding to the 5 newly generated ERα ChIP-seq datasets have been deposited at the European Genome-phenome Archive (EGA) (54) – which is hosted by the EBI and the CRG – under accession number EGAS50000000008, while processed data can be found on GEO under accession number GSE244845 (ChIP-seq subseries: GSE244840). Called peaks of ERα ChIP-seq for MCF7 (GSM798423, GSM631484, GSM1967545), T-47D (GSE68359, GSE32222) (6,13) and ZR-75-1 (GSE25710) are available on GEO.

Hi-C raw and processed data can be found in GEO under accession number GSE244845 (Hi-C subseries: GSE244844).

STARR-seq raw data are available at EGA’s accession number EGAS50000000009, while processed data are included in Supplementary Table 5.

The mass spectrometry proteomics data have been deposited to the ProteomeXchange Consortium (55) via the PRIDE (56) partner repository with the dataset identifier PXD045526.

## AUTHOR CONTRIBUTIONS

Design of the study: S.E.P.J., W.Z. Performance of experiments: S.E.P.J., S.G., S.S., E.Y., C.F.H., M.D.C. Data analysis: S.E.P.J., S.G., T.M., B.A., Y.K., S.C. Wrote the manuscript: S.E.P.J., W.Z. Project supervision: G.K., N.L., M.V., S.C.L., W.Z. Critically read and edited the manuscript: S.G., G.K., N.L., S.S., M.D.C, G.K., N.L., M.V., S.C.L., W.Z.

All authors reviewed and approved the manuscript prior to submission.

## ACKNOWLEDGEMENTS

The authors would like to thank patients who donated tumor material for scientific research and all researchers involved in the generation and availability of the public datasets used in this manuscript.

## FUNDING

This project was funded by Alpe d’HuZes / Dutch Cancer Society (NKI-2014-7140). The Vermeulen and Zwart lab are part of the Oncode Institute, which is partly funded by the Dutch Cancer Society. Per BCAC terms of use, we emphasize their analyses were supported by the Government of Canada through Genome Canada and the Canadian Institutes of Health Research, the “Ministère de l’Économie, de la Science et de l’Innovation du Québec” through Genome Québec and grant PSR-SIIRI-701, The National Institutes of Health (U19 CA148065, X01HG007492), Cancer Research UK (C1287/A10118, C1287/A16563, C1287/A10710) and The European Union (HEALTH-F2-2009-223175 and H2020 633784 and 634935).

## Conflict of interest statement

None declared.

## Notes

### Competing Interest Statement

The authors have declared no competing interest.

